# Cognate T and B cell interaction and association of Follicular helper T cells with B cell responses in *Vibrio cholerae* O1 infected Bangladeshi adults

**DOI:** 10.1101/306365

**Authors:** Rasheduzzaman Rashu, Taufiqur Rahman Bhuiyan, Mohammad Rubel Hoq, Lazina Hossain, Anik Paul, Ashraful Islam Khan, Fahima Chowdhury, Jason B Harris, Edward T Ryan, Stephen B Calderwood, Ana A Weil, Firdausi Qadri

## Abstract

*Vibrio cholerae* O1 can cause life threatening diarrheal disease if left untreated. A long lasting immune response, producing 3-5 years of protection from subsequent, symptomatic disease following natural infection, is mediated by B cell mediated humoral immunity. T cells can play critical roles in inducing such immunity. However, the mechanism of T cell dependent B cell maturation and whether a key sub-population of T cells are involved is not well established in cholera. We hypothesized that a specific population of T cells, follicular helper T (Tfh) cells, are involved in B cell maturation following cholera; we used flow cytometry, culture and colorimetric assays to address this question. We found that *V. cholerae* infection induces significant increase in circulating Tfh cells expressing B cell maturation associated protein CD40L early in disease. The increased Tfh cells expressing CD40L recognize cholera toxin most prominently, with lessened responses to two antigens tested, *V. cholerae* membrane preparation (MP) and *Vibrio cholerae* cytolysin (VCC). We further showed that early induction of Tfh cells and CD40L was associated with later memory B cell responses to same antigens. Lastly, we demonstrated *in vitro* that Tfh cells isolated after cholera can stimulate class switching of cocultured, isolated B cells from patients with cholera, leading to production of the more durable IgG antibody isotype. These studies were conducted on circulating Tfh cells; future studies will be directed at examining role of Tfh cells during cholera directly in the gut mucosa of biopsied samples, at the single cell level if feasible.

## Introduction

Cholera is a life threatening diarrheal disease. It is estimated that 3-5 million cases and 100,000 deaths occur globally every year (1). *Vibrio cholerae* O1, the causative agent of cholera, produces cholera toxin (CT), *Vibrio cholerae* cytolysin (VCC), a variety of membrane-associated proteins and the toxin-coregulated pilus A (TcpA), all involved in immunogenicity and pathogenicity (2, 3). CT, which is a T cell dependent protein antigen, induces salt and water loss in the intestine, the major cause of dehydration and death (4). Natural infection induces long-lasting CT-specific IgG producing memory B cells (4). Such T cell dependent protein antigens can induce anamnestic memory B cell responses on re-exposure, while T cell independent antigens like lipopolysaccharide (LPS) and the O-specific polysaccharide it contains fail to produce such durable responses after *V. cholerae* infection (4). In addition to memory B cells that develop to cholera protein antigens after infection, memory helper T cell responses to protein antigens also develop after cholera by day 7, prior to initiation of memory B cell responses to the same antigens; memory helper T cell responses are not seen to LPS (3, 4).

Among the various types of T cells, Tfh are a subpopulation of CD4+ T cells that are found in the secondary lymphoid organs as well as peripheral blood (5), express CXCR5 on their cell surface and, following stimulation by cognate antigen, migrate into the B cell zones of lymphoid organs, mediated through CXCR5-CXCL13 crosstalk (6, 7). There, Tfh cells interact with B cells that present the cognate antigen, with the help of MHCII on their cell surface to be recognized by the TCR; this recognition is facilitated by the interaction of CD40L on the surface of the activated Tfh and CD40 on the surface of the antigen-presenting B cell (8). This contact dependent interaction, plus the secretion of cytokines, including IL-21 and IL-4 by the Tfh cell, helps to trigger the formation of germinal centers. The CD40L-CD40 interaction is essential not only for formation of the germinal center but also for its maintenance in the secondary lymphoid organs (9). In addition, this bi-directional interaction of Tfh-germinal center (GC) B cells facilitates the survival of those specific GC B cells. The Tfh-B cell interaction then further leads to B cell maturation, including isotype switching and somatic hypermutation by inducing activation-induced cytidine deaminase in B cells; this leads to either the production of mature plasma cells secreting high affinity, antigen-specific antibodies or the production of antigen-specific memory B cells (10, 11). The role of Tfh cells, including expression of co-stimulatory molecules, has not been studied in cholera patients, and whether this same interaction leads to B cell activation and maturation in this mucosal infection is not known. We performed the current study to investigate these potential interactions.

## Materials And Methods

### Study subjects and overview

Patients with symptoms of severe dehydration were admitted to the International Centre for Diarrheal Diseases Research, Bangladesh (icddr,b), Dhaka hospital. Patients’ stool was collected, and *V. cholerae* O1 infection was confirmed by dark field microscopy and by culture on selective taurocholate-tellurite gelatin agar media described elsewhere (12). The serogroup and serotype of the infecting *V. cholerae* O1 strains were determined by agglutination test by anti-O1, anti-Ogawa and anti-Inaba specific monoclonal antibodies (13, 14). Patients confirmed to have *V. cholerae* O1 infection (n=34) were enrolled following informed consent, and blood was collected acutely during infection (2^nd^ day of hospitalization), during early convalescence (7^th^ day after hospitalization) and late in the convalescent period (30^th^ day after hospitalization). Patients were excluded from this study if coinfected with other enteric pathogens or parasites. The study was approved by Ethical and Research Review Committees of the International Centre for Diarrhoeal Disease Research, Dhaka, Bangladesh (icddr,b) and the Institutional Review Board of Massachusetts General Hospital.

### Separation of peripheral blood mononuclear cells (PBMCs) and detection of CD40L expressing Follicular helper T cells

Heparinized venous blood was collected in BD vacutainer tubes and diluted with an equal volume of phosphate buffered saline (PBS, 10 mM, pH ~7.2). Diluted blood was carefully loaded on Ficoll-Isopaque (Pharmacia, Uppsala, Sweden) without disturbing the Ficoll layer, and then centrifuged at 700 g for 30 minutes at 20°C. After centrifugation, PBMCs were collected from the top of the Ficoll layer, washed and counted with a haemocytometer, and cell viability was assessed using trypan blue dye (15). Separated PBMCs were incubated with anti CD3-PE Texas Red (Invitrogen, CA), anti CD4-Amcyan, anti CXCR5-Alexa-Fluor-488, and anti CD40L-APC-eFluor700 (BD Bioscience, San Jose, CA) fluorochrome-tagged antibodies. After 45 min of incubation in the dark at 4°C, extra fluorochrome was washed out with PBS supplemented with 2% fetal bovine serum (FBS) (HyClone, Logan, UT), the labeled cells diluted to the desired volume and then cell counts acquired with a FACSAriaIII instrument (BD Bioscience, San Jose, CA) and the FACSDiva software program. We analyzed the acquired data with FlowJo software (TreeStar, Inc., OR), separating the lymphocyte population depending on side and forward scattered light. Lymphocytes were then identified by gating on the CD3-positive population, followed by the CD4-positive population within the CD3-positive cells. Tfh cells were selected by CXCR5 expression on the cell surface of CD4-positive cells, and the Tfh population was selected using CD40L expression. The results are expressed as percentages.

### Antigens used for short-term culture and CD40L expression, memory B cell enumeration and depleted Tfh cell and B cell co-culture

Membrane preparation (MP), *V. cholerae* cytolysin (VCC), and cholera holotoxin containing the G33D variant of the B subunit (mCT) were used in this study. VCC (a gift from Kalyan K. Banerjee) monomer was produced from a nonclinical *V. cholerae* O1 strain that is biochemically and immunologically similar to *V. cholerae* O1 (16). MP was prepared from *V. cholerae* O1 El Tor strain N16961 grown in AKI medium and the most abundant proteins were characterized by mass spectrometry (3, 17). mCT (a gift from Randall K Holmes) has less binding affinity to eukaryotic cell surface ganglioside (GM1) due to an aspartic acid substitution by glycine on the B subunit, and is a less toxic variant of wild type cholera toxin (CT), useful in cell culture conditions (18). VCC was used at a concentration of 2.5ng/ml. MP and G33D mCT were used at a concentration of 10μg/ml. We also used phytohaemagglutinin (PHA) (Remel, USA) as a positive control at 1μg/ml and media only as a negative control in cell stimulations in culture. In addition, recombinant cholera toxin B subunit rCTB (a gift from Ann-Mari Svennerholm) was used for enumeration of antigen-specific memory B cells in the peripheral blood and in enzyme-linked immunosorbent assay (ELISA). Staphylococcal Enterotoxin B from *Staphylococcus aureus* (SEB) (Sigma, USA) was used as a positive control for co-culture of depleted Tfh cells and B cells, as done previously (5, 19).

### Whole blood stimulation by FASCIA

We used Flow cytometric Assay of Specific Cell-mediated Immune responses in the Activated whole blood (FASCIA) to measure cellular proliferation in response to antigenic stimulation as previously described (3, 20, 21). In brief, we collected whole blood in a lithium-heparinized vacutainer tube and diluted it to a ratio of 1:8 in Dulbecco modified Eagle medium (Gibco, NY) supplemented with 1% gentamicin, 1% mercaptoethanol, and 10% heat-inactivated fetal calf serum. We added 100ul of stimulating antigen, control antigen, or additional medium to each tube of 400ul of diluted blood in a 5-ml polystyrene tube. After 6 hours of *in vitro* stimulation in a humidified atmosphere with 5% CO_2_ at 37°C, cells were centrifuged and the supernatant was discarded. Whole blood was stained with anti-CD3-PE Texas Red (Invitrogen, CA), anti-CD4-Amcyan, anti-CXCR5-AF488, and anti-CD40L-APC eFluor700, monoclonal antibodies (BD Bioscience, San Jose, CA). Red blood cells were lysed with ammonium chloride solution containing potassium chloride and EDTA, and the remaining cells were centrifuged, washed and fixed with BD stabilizing fixative (BD Bioscience, San Jose, CA). Cell counts were acquired for 4 min by FACS Aria III and FACS Diva software (BD Bioscience). Acquired cell counts were analyzed, counted by FlowJo (San Jose, CA). The magnitude of stimulation is expressed as the ratio of lymphocyte count expressing a certain protein on their cell surface with antigenic stimulation to the count without stimulation (21, 22). The ratio is referred to as a stimulation index (SI). An SI value equal to “1” indicates that stimulation is equal in samples with or without a *V. cholerae* antigen, and more than “1” indicates *V. cholerae* antigen-specific stimulation. Lymphocytes were gated and counted depending on forward and side scattering characteristics. CD3+ cells were gated from lymphocytes and CD4+ cells were gated from CD3+ cells. Tfh cells were counted from CD4+ cells based on surface expression of CXCR5 on cell surface. CD40L expressing cells from the Tfh cell pool were analyzed.

### Enzyme-linked immunosorbent spot (ELISPOT) assay to measure memory B cells

Memory B cells were measured by the ELISPOT assay described previously (23, 24); this assay is optimized for stimulating differentiation of memory B cells into terminal, spot producing antibody secreting cells (ASCs). In brief, PBMCs were seeded (5×10^5^ cells/well) in cell culture plates (BD Biosciences, San Jose, CA) with RPMI 1640 (Gibco, Carlsbad, CA) and 10% fetal bovine serum (FBS) (HyClone, Logan, UT). Cells were stimulated with a cocktail of B cell mitogens containing 6 μg/ml CpG oligonucleotide (Operon, Huntsville, AL), a 1/100,000 dilution of crude pokeweed mitogen extract, and a 1/10,000 dilution of fixed *Staphylococcus aureus* Cowan (Sigma, St. Louis, MO). Plates were incubated at 37°C in 5% CO_2_ for 6 days. The cells were harvested, washed and was transferred onto nitrocellulose membrane-bottom plates (MSHAN-4550; Millipore, Bedford, MA) for 5 hours. The plates were coated previously with GM1 ganglioside (3 nmol/ml) overnight followed by recombinant CTB (2.5 μg/ml), or with 5 μg/ml affinity-purified goat anti-human immunoglobulin (Jackson Immunology Research, West Grove, PA), or 2.5 μg/ml keyhole limpet hemocyanin as a negative control (Pierce Biotechnology, Rockford, IL). Each coating was followed by blocking with RPMI 1640 containing 10% FBS. After incubation, cells were washed out and horseradish peroxidase-conjugated goat anti-human IgA (Southern Biotech, USA) and alkaline phosphatase conjugated goat anti-human IgG (Southern Biotech, USA) were added. Following an overnight incubation at 4°C, plates were developed with 3-amino-9-ethyl carbazole and 5-bromo-4-chloro-3-indolyl-phosphate/Nitro blue tetrazolium for developing IgA and IgG spots. The spots were counted and results were expressed as the percentage of antigen-specific memory B cells out of the total isotype-specific memory B cells.

### Co-culture of B cells and T follicular helper cells and ELISA measurement of total IgG production after co-culture

We largely followed a previously published procedure (5), with the exception that the co-cultures contained *V. cholerae* antigens for Tfh stimulation, with SEB as a positive control and media only as a negative control. PBMCs were separated by Ficoll-isopaque at day 7 after hospitalization and stained with anti-CD3-PE Texas Red, anti-CD4 Amcyan, anti-CD45R0 PE, anti-CXCR5-Alexa Flour 488, and anti-CD19-PerCP-Cy 5.5 and kept in the dark for 45 min at 4ºC. Then, the cells were washed with PBS 2% FBS to remove any unbound fluorochrome tagged antibody, passed through a 40μm cell strainer (BD) and diluted with PBS supplemented with 2% FBS to the desired concentration for sorting approximately 10 million cells per milliliter of buffer. CD19+ B cells, CD3+CD4+CD45R0+CXCR5+ memory Tfh cells (Tfh+) and CD3+CD4+CD45R0+CXCR5-memory cells (Tfh-) were sorted with a FACS AriaIII sorter and collected in three different 1% BSA pre-coated 5 ml polystyrene tubes (BD Falcon). After sorting, the separated cells were centrifuged and diluted to the desired volume. Sorted cell types were rerun to check the purity of the sorted cells; the percentage of purity was more than 98 percent. Sorted B cells (50,000 cells) were co-cultured with Tfh+ (50,000 cells) or Tfh-(50,000 cells) cells with antigenic stimulation for seven days in a humidified 5% CO_2_ incubator at 37°C. After seven days of co-culture, culture supernatants were collected and stored at −80°C to measure secreted IgG.

ELISA was used to measure the total secreted IgG from co-cultured supernatants (23). In brief, a 96-well polystyrene plate (Nunc F) was coated with 5 μg/ml affinity-purified goat antihuman immunoglobulin (Jackson Immunology Research, West Grove, PA). Culture supernatant or ChromPure Human IgG molecule (Jackson Immunology Research, West Grove, PA) was applied and incubated for 90 minutes, followed by the addition of horseradish peroxidase-conjugated secondary antibodies to human IgG (Jackson Immunoresearch, USA). The plate was developed with ortho-phenylene diamine (Sigma, St. Louis, MO) in 0.1 M sodium citrate buffer (pH 4.5) and 0.1% hydrogen peroxide. The developed color was read using a microplate reader (Eon). Sample concentrations were determined by comparing to the known concentrations of pure IgG.

### Statistical analysis

We used Graphpad Prism 5.0 for statistical analyses and to generate figures. We compared data using paired t-tests; the Wilcoxon signed-rank test was used to measure the responses of each patient on different days. We used Pearson’s correlation to assess the relationship between T and B cell responses. All reported *P* values are two tailed with 95% confidence intervals. *P*≤ 0.05 is considered as statistically significant.

## Results

### Study population

A total of 34 adult patients were recruited in this study. Demographic and clinical features of these study patients by age, sex, blood group and duration of diarrhea at presentation are shown in Table 1. Sample size in each experiment varied and was restricted by the allowed small volume of blood obtained from each participant. The enrollment of cholera patients in each group followed the progression of the experiments and generation of data, which then determined the investigators decision for the next experimental step. Twenty samples were analyzed for the determination of frequency of circulating Tfh cells and CD40L expressing Tfh cells after infection (Fig. 1), 18 were analyzed for evaluation of antigen-specific memory T cell responses (Fig. 2), 10 were used for the correlation analysis (Fig. 3) and 11 for the depletion and co-culture experiments (Fig. 4).

**Table 1.** Demographic and clinical features of patients participating in this study.

| Characteristics | Patients |
| --- | --- |
| Number of study participants | 34 |
| Median age in years (25<sup>th</sup>, 75<sup>th</sup> percentile) | 35 (29.8, 40.3) |
| Gender, female (%) | 12 (35.3) |
| Mean (SD) duration of diarrhea, hours | 30.7 (56.1) |
| Blood group (%) |  |
| A | 25.8 |
| B | 45.2 |
| O | 25.8 |
| AB | 3.2 |

**Fig 1.**
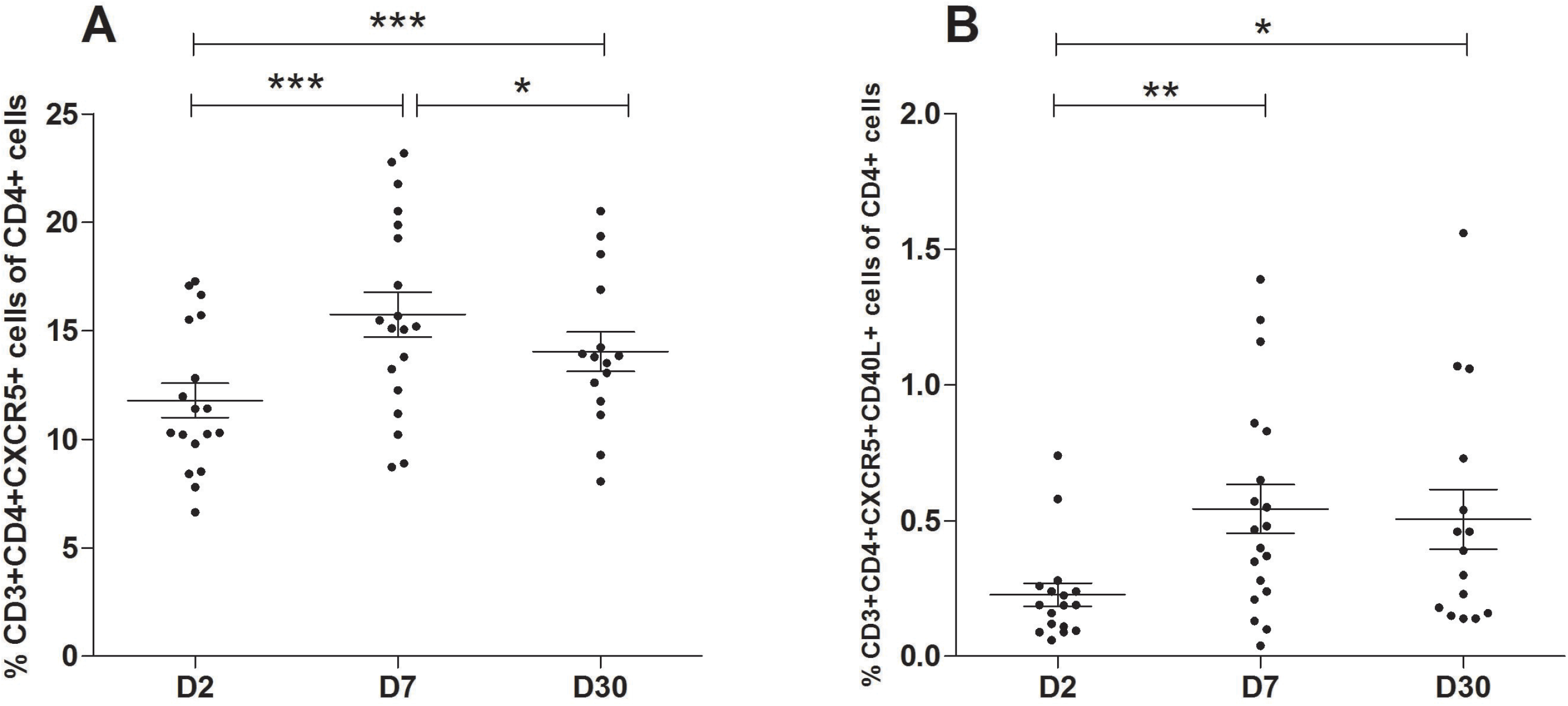
(A) Frequency of Tfh cells (CD3+CD4+CXCR5+) and (B) frequency of CD40L-expressing Tfh cells (CD3+CD4+CXCR5+CD40L+) plotted as a percentage of total CD4+ cells in peripheral blood at day 2 (D2), day 7 (D7) and day 30 (D30) following onset of cholera (n=20). Data are presented as the mean percentage of Tfh cells and the CD40L+ Tfh cells, with error bars representing standard error of the mean. An asterisk represents a significant increase from day 2 or decrease from day 7 (* <0.05, **<0.01, ***<0.001) using a paired t-test.

**Fig 2.**
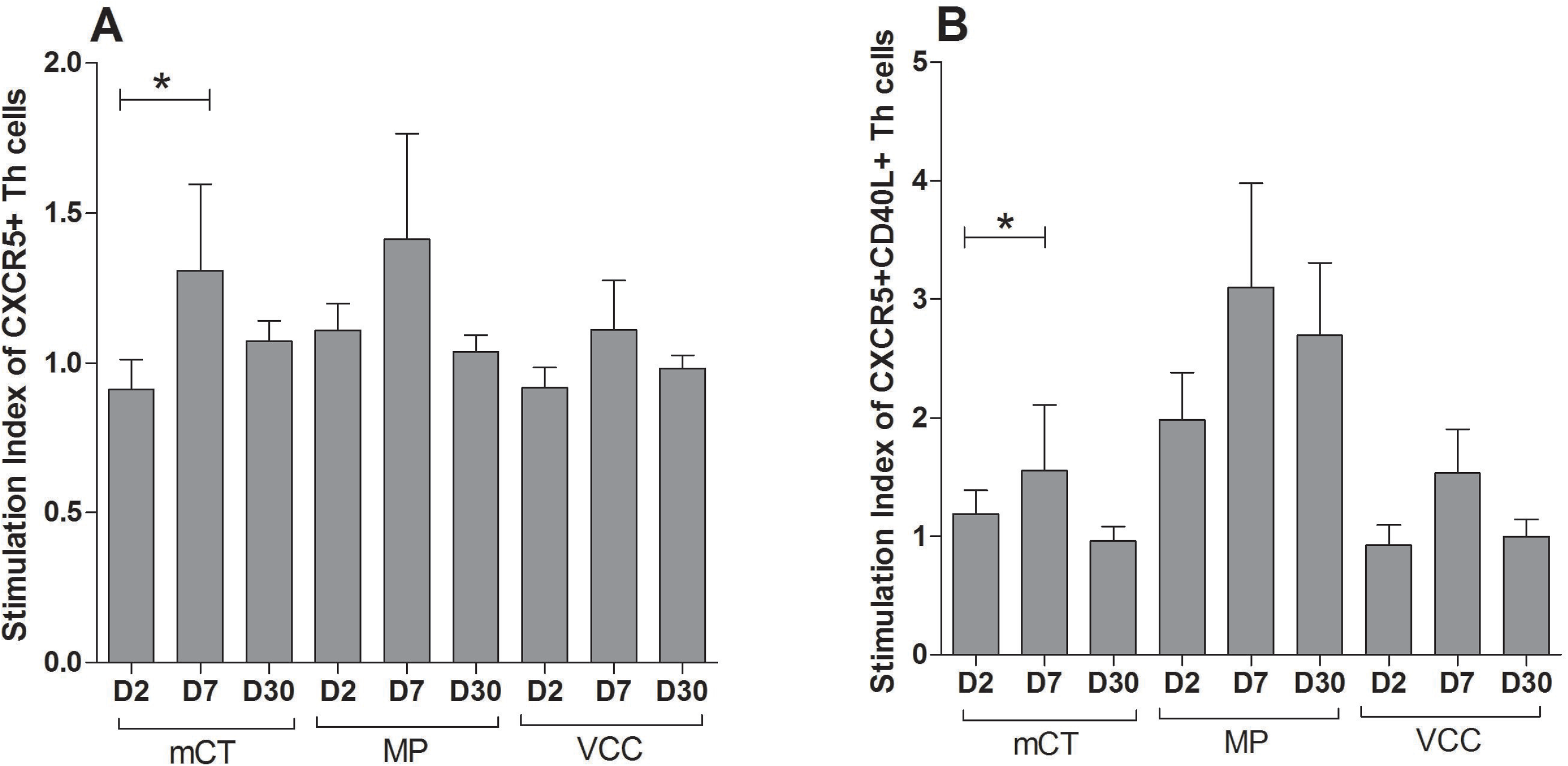
Evaluation of antigen-specific proliferation (n=18) of Tfh cells (A) and CD40L-expressing Tfh cells (B) after cholera; cells were stimulated with G33D mutant cholera toxin (mCT), *V. cholerae* membrane preparation (MP), and *V. cholerae* cytolysin (VCC). Data are presented as mean stimulation indices (SI) of Tfh cells and CD40L expressing Tfh cells, with error bars representing standard errors of the mean. Asterisks represent a P-value less than or equal to 0.05 and represent a significant increase from day 2.

**Fig 3.**
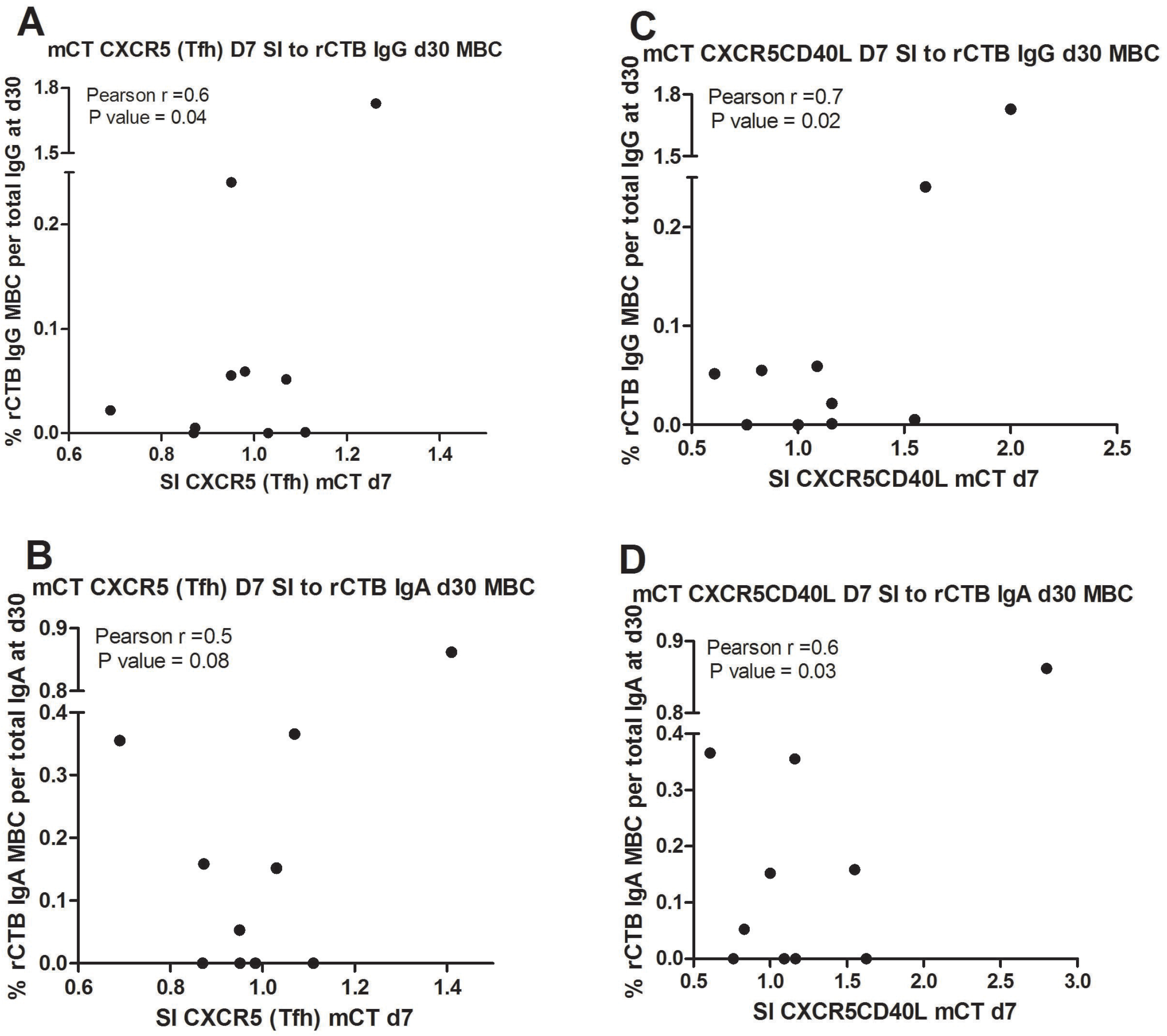
Correlation of day 7 circulating antigen-specific Tfh and CD40L-expressing Tfh cells with subsequent antigen-specific memory B cells at day 30 in the same patients (n=10). The stimulation index (SI) of Tfh cells (CXCR5+) or CD40L-expressing Tfh cells were analyzed for correlation with rCTB-specific IgG memory B cells (A, B), and with rCTB-specific IgA memory B cells (C, D). Correlation was determined by the Pearson correlation coefficient.

**Fig 4.**
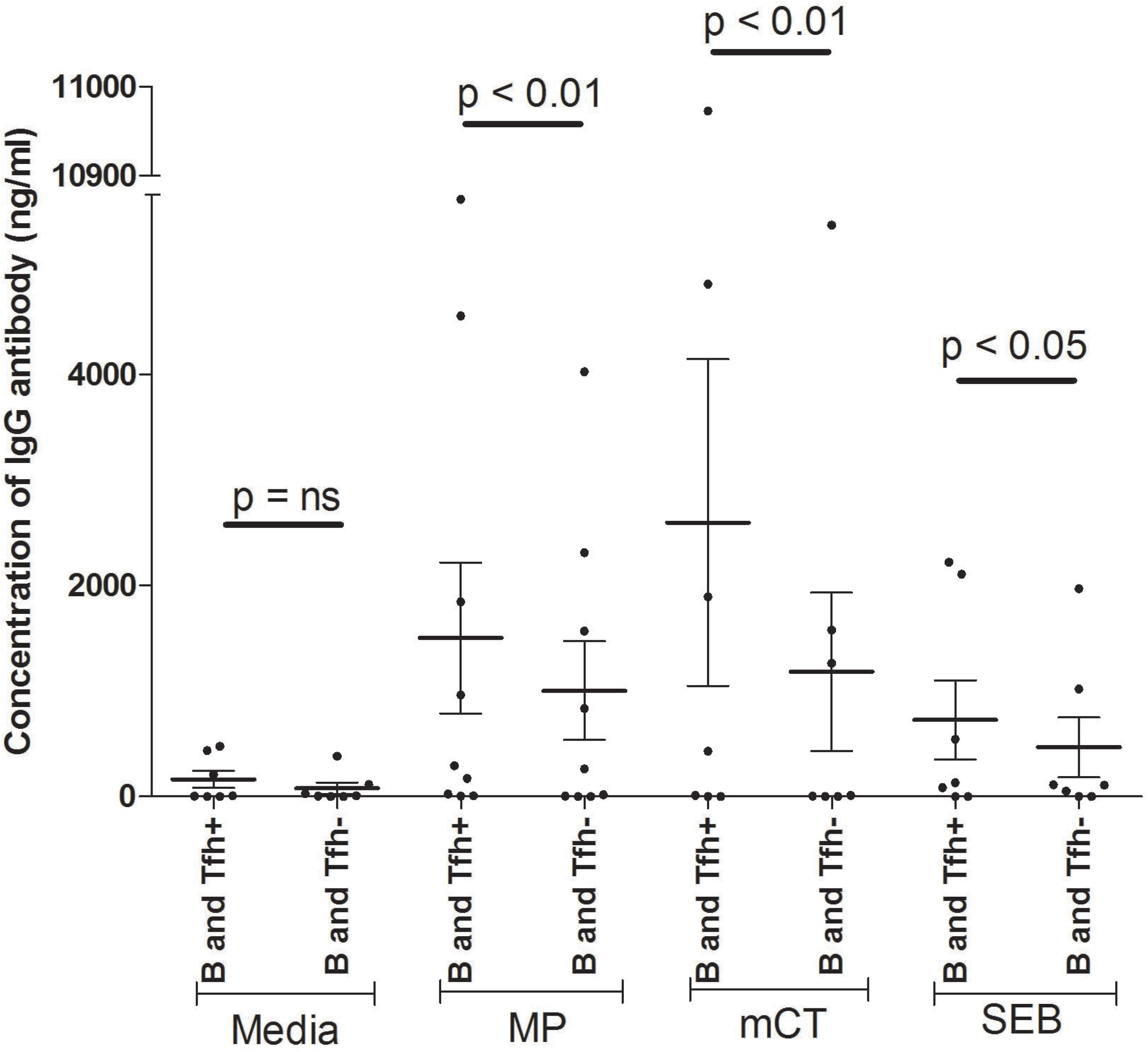
*In vitro* co-culture of Tfh cells or Tfh depleted T helper cells with B cells after 7 days, with antigenic stimulation (MP, mCT, or SEB) and a media only control (n=11). Data are presented as the mean concentration of IgG in ng/ml of culture supernatant, with error bars representing the standard errors of the mean. B, Tfh+ and Tfh-cells are defined as CD19+ cells, CD3+CD4+CD45R0+CXCR5+ cells and CD3+CD4+CD45R0+CXCR5-cells, respectively. An asterisk represents a significant decrease of IgG after depletion of Tfh compared to nondepleted Tfh cells (* <0.05, **<0.01).

### Frequency of Tfh cells and CD40L expressing Tfh cells after onset of cholera

Whole blood was diluted and stained with anti-CD3, -CD4, -CXCR5 and -CD40L antibodies, and the resulting populations were counted by flow cytometry. The percentage of Tfh cells in peripheral blood was significantly increased at day 7 (15.3±1.1) following *V. cholerae* infection compared to day 2 (11.5±0.8) (*P*<0.0001) and waning by day 30 (13.8±0.9); however, the level remained elevated above baseline (Fig. 1A). CD40L-expressing Tfh cells were also significantly elevated at day 7 (*P*=0.01) and day 30 (*P*=0.05) following infection compared to day 2 (Fig. 1B). The expression of CD40L-expressing Tfh at day 30 did not decrease compared to day 7.

### Antigen-specific Tfh cell proliferation and B cell maturity associated protein CD40L expression

Whole blood was obtained on day 2, day 7 and day 30 after infection, and diluted and stimulated with *V. cholerae* antigens: mCT, MP, VCC, or unstimulated for 6 hours. The Stimulation Index (SI) (the ratio of cell number after stimulation with an antigen to cell number without stimulation) was calculated for each antigen. mCT-specific Tfh cells were significantly increased at day 7 after infection compared to day 2 (*P*=0.04, Fig. 2). mCT-specific Tfh cells that express CD40L were also significantly elevated following infection (*P*=0.02). The SI for MP and VCC antigen showed similar trends but did not reach statistical significance.

### Antigen-specific Tfh cells and CD40L-expressing Tfh cells in the circulation on day 7 are correlated with subsequent antigen-specific IgA and IgG producing memory B cells on day 30 following cholera

mCT-specific Tfh and CD40L expressing Tfh cells were determined by FASCIA on day 7 of infection as above, and rCTB-specific IgA and IgG-secreting memory B cells were evaluated by ELISPOT on day 30 following infection in the same patients, and the results examined for correlation. Tfh cells specific for mCT on day 7 following infection correlated with IgG secreting memory B cells specific for rCTB on day 30 (Pearson r= 0. 6, *P*=0.04, Fig. 3); a similar trend that did not reach statistical significance was seen for IgA secreting memory B cells on day 30. mCT-specific CD40L expressing Tfh cells on day 7 also correlated with IgA (Pearson r= 0.7, *P*=0.02) and IgG (Pearson r= 0.7, *P*=0.01) secreting rCTB-specific memory B cells on day 30 following infection.

### Depletion of memory Tfh cells reduces the amount of IgG secretion by autologous B cells into culture supernatant following an *in vitro* co-culture of cells recovered from peripheral blood of cholera patients

B cells and Tfh cells were sorted from the peripheral blood of cholera patients (n=11) by flow cytometry. Sorted B cells (CD19+ cells) were cultured with sorted memory Tfh cells (CD3+CD4+CD45R0+CXCR5+ cells) with antigenic stimulation; these cells secreted a significant amount of IgG in the presence of two *V. cholerae* antigens, MP or mCT, compared to B cells co-cultured with a fraction depleted in Tfh cells (CD3+CD4+CD45R0+CXCR5-cells) (Fig. 4). In comparison, without antigenic stimulation, B cells did not produce a significant amount of IgG in the presence of Tfh cells compared to depleted Tfh cells.

## Discussion

Infection with *V. cholerae* produces protective immunity and induces antigen specific IgG antibodies and IgG memory B cells to protein antigens such as CT that persist at least one year after *V. cholerae* infection (4, 25). In addition, an oral cholera vaccine containing added rCTB has been shown to elicit an antigen-specific IgG-immune response to CTB comparable to natural infection (23). Both natural infection and existing vaccines with protein components induce B cells to produce and secrete durable, class-switched IgG antigen-specific antibody. However, these durable and avid class-switched antibodies are only seen in response to protein antigens and not observed for a non-protein antigen such as lipopolysaccharide (15). This suggests that following *V. cholerae* infection, maturation of a repertoire of B cells specific for T cell dependent antigens relies on interaction with a certain population of helper T cells (26). However, the mechanism of this T cell dependent B cell maturation in a mucosal infection such as cholera is not fully defined. Here, we have shown that natural infection with *V. cholerae* induces antigen-specific Tfh cell proliferation and expression of CD40L, and that these Tfh cells then provide antigen-specific help for B cell maturation, class switched antibody secretion and subsequent development of antigen-specific memory B cell responses to the same antigens in peripheral blood. Lastly, in an *in vitro* system, B cells co-cultured with Tfh cells recovered from cholera patients resulted in increased production of class switched IgG antibodies by the B cells in culture. This suggests that Tfh cells are important in regulating B cell proliferation and maturation in this mucosal infection, leading to class switching and likely somatic hypermutation, as seen in other infections (24, 27–30).

In normal physiologic conditions, naïve T cells migrate from the thymus to the CCL19 - and CCL21- rich region of a secondary lymphoid organ utilizing chemokine receptor CCR7, where they surveil for foreign antigens (31). After interacting with antigen presenting cells, these naïve T cells become a heterogeneous population of antigen primed T cells that includes CXCR5 - expressing follicular helper T cells (Tfh), which migrate to the B cell zone of the lymphoid tissue utilizing interaction with CXCL13 (32, 33). The proper spatial and temporal contact-dependent interaction between an antigen-primed Tfh cell expressing CD40L and CD40 expressing B cells in the T cell-B cell border of a secondary lymphoid organ is necessary for maturation of those B cells, including class switching of antibody production (10, 32, 34, 35). Without proper help either by direct contact and/or cytokine production from Tfh cells, B cells do not undergo either class switching or somatic hypermutation to produce higher avidity IgA and IgG isotype antibodies (36, 37).

Whether similar events occur in lymphoid tissues of gastrointestinal mucosa or the draining lymph nodes following mucosal infection is not well defined. Here, we assayed peripheral blood of cholera patients in Bangladesh for circulating Tfh cells and their subsequent effect on antigen-specific B cell events following a mucosal infection caused by *V. cholerae*. Measuring these events in the circulation assumes that a small portion of the antigen-specific Tfh cells stimulated and active in germinal center follicles in lymphoid tissues migrate into, and can be assayed in peripheral blood, as shown by others (5, 19, 38). We found that Tfh cells are upregulated in the circulation on day 7 after *V. cholerae* infection, and express higher amounts of the surface B cell maturation associated marker CD40L. This suggests that similar events are taking place in the germinal centers (GC) of secondary lymphoid tissues after cholera. Previous groups have shown that CD40L is critical for GC development and maintenance (39), proliferation and maintenance of highly proapoptotic B cells within GC (9), and for plasma cell formation (40). We have previously shown that CT-specific plasmablasts also peak in the circulation at day 7 after infection, at the same time as the circulating (and presumably lymphoid tissue-associated) Tfh cells expressing CD40L peak (41). The fraction of CD40L expressing Tfh cells in the circulation may underestimate the fraction of these cells in GC of secondary lymphoid tissues. However, obtaining direct samples of secondary lymphoid tissues from cholera-infected patients *in vivo* to examine this correlation is challenging from a human study standpoint.

We demonstrated that the expanded, circulating Tfh cells at day 7 after infection with increased expression of CD40L are *V. cholerae* antigen-specific, particularly to the highly immunogenic G33D mutant of cholera toxin. We have previously shown a more durable memory B cell response to CT than to the T cell independent antigen LPS, suggesting that the interaction of Tfh cells expressing CD40L with cognate B cells orchestrates this more durable memory B response to CT in cholera patients (4). A membrane preparation (MP) of *V. cholerae* was less effective in inducing Tfh cell proliferation and CD40L expression compared to mCT, which is consistent with our previous findings (22). One possible explanation for this might be the heterogeneous mixture of proteins in MP, such that the quantity of each individual protein might not overcome a threshold level of T cell interaction with cognate B cells. This might be addressed by purifying and testing the more abundant proteins from MP, such as outer membrane protein U (OmpU). Although VCC has been previously shown to be immunogenic following cholera (3), we found here that VCC was less effective at stimulating Tfh cell proliferation and CD40L expression. A mutant form of VCC with lessened cytotoxicity may better assess the immunogenicity of VCC in this cellular proliferation assay.

A previous study showed that naïve T cells take approximately 3 to 4 days to enter the B cell zone of lymphoid tissues following expression of CXCR5 on their surface starting at 36 hours (42); naïve T cells need to accumulate sufficient CXCR5 to overcome signals from CCR7 that retain these cells in the T cell zone (32). Therefore, most antigen-activated Tfh cells are in proximity to cognate B cells in the B cell region of GC around day 7 following infection. This timing is consistent with our previous findings that the T cell response on day 7 after cholera vaccination correlates with later B cell events in vaccine recipients (22).

A recent elegant study by Cardeno *et al*., published while this manuscript was being finalized, examined circulating Tfh cells in adults following oral vaccination with an inactivated ETEC vaccine (19), and the relationship of these circulating Tfh cells and B cell events. Although they found a small increase in overall CXCR5-expressing Tfh cells in the circulation 7 days after vaccination, they found a larger increase in the fraction of these Tfh cells expressing the activation marker ICOS; this is similar to our results with CD40L expression on circulating Tfh cells after cholera. Cardeno *et al*. (19) showed that circulating Tfh cells after ETEC vaccination express increased amounts of PD1 and β7, and produce increased IL-21 after stimulation. ETEC vaccination also led to an increased number of plasmablasts in the circulation on day 7 after vaccination, and the majority of these expressed IgA and β7, consistent with mucosal homing. In this study (19), there was a correlation of the fold rise of circulating activated Tfh cells in the circulation on day 7 and the fold rise in circulating IgA plasmablasts on day 7.

To directly model the effect of the circulating, antigen-specific Tfh cells on B cell maturation in our study, we utilized short term *in vitro* co-culture of Tfh cells and isolated B cells from cholera patients 7 days following infection. We showed that co-culture of CXCR5+, CD40L expressing Tfh cells with autologous B cells in the presence of *V. cholerae* antigens stimulated production of IgG antibody in culture supernatants, while depletion of the Tfh cells from the pool of circulating T cells resulted in a significantly lower amount of IgG antibody production. A previous study by another group utilized co-culture of circulating B cells with CXCR5+ T cells from patients with the autoimmune disease juvenile dermatomyositis (5) and showed increased production of IgG and IgA antibodies in culture supernatants compared to coculture of B cells with CXCR5-depleted T cells. Cardeno *et al*. (19) similarly isolated circulating Tfh cells and B cells following ETEC vaccination and showed that *in vitro* co-culture produced an increased frequency of IgA secreting plasmablasts differentiated from the B cells in the culture, increased amounts of total IgA and IgG secreted into culture supernatants, and increased IgA in culture supernatant specific for the B subunit of ETEC heat-labile enterotoxin from the vaccine. Of note, the two previous studies (5, 19) utilized the superantigen SEB in the co-culture experiments to stimulate the Tfh cells *in vitro*. We also used SEB as a positive control, but also showed that cholera-specific antigens could be utilized for antigenic stimulation as well, as opposed to a superantigen.

Cardeno *et al*. (19) also examined patients undergoing ETEC revaccination 1-2 years later to show a correlation between the fold rise of ICOS+, circulating Tfh cells on day 7 post primary vaccination, and the fold rise of IgA responses in circulating, mucosal antibody-secreting cells to ETEC antigens following revaccination 1-2 years later. We similarly found a correlation between CD40L expressing Tfh cells circulating on day 7 following cholera, and IgA and IgG memory B cell responses to rCTB on day 30 following infection. Both these observations are consistent with a model whereby Tfh cells in gut-associated lymphoid tissues induce B cells to mature to class switched IgA and IgG antibody producing plasma cells, as well as IgA and IgG memory B cells that are then found in the circulation following cholera (23).

Recently, Holmgren *et al*. summarized some novel approaches to correlate early human mucosal immune responses with subsequent protection following mucosal vaccination, which include the investigation of gut-originating antibody-secreting cells (ASCs), memory B cells and Tfh cells from samples of peripheral blood during their recirculation from lymphoid tissues (43). A previous report, however, suggests that in the absence of cognate interactions, the function of CD4 Tfh cells in germinal centers of Peyer patches, as opposed to other lymphoid tissues, is not clear (44). To better understand the role of Tfh cells in immune responses in cholera patients, further investigation including single cell studies from gut biopsies at the site of *V. cholerae* replication may be needed.

Our study and the previous work of others (5, 19) are consistent with a model in which activated Tfh cells in lymphoid tissues interact with B cells following mucosal infection or vaccination to stimulate affinity maturation and differentiation of antigen-specific plasmablasts and memory B cells, and that these events are important in development of mucosal memory to T cell dependent antigens. However, the protective immune response following cholera appears to be against the the O-specific polysaccharide component of LPS which is the immunogenic part (45); how mucosal memory develops to this T cell independent antigen is not yet clear, and also requires further study.

## Acknowledgement

This work was supported by the International Centre for Diarrhoeal Disease Research, Bangladesh (icddr,b). This study was also supported by grants from the National Institutes of Health, including grants from the National Institute of Allergy and Infectious Diseases (AI106878 [E.T.R. and F.Q.] and AI058935 [S.B.C., E.T.R., and F.Q.], AI103055 [J.B.H. and F.Q] and K08 AI123494 [A.A.W.], as well as the Fogarty International Center (Training Grant in Vaccine Development and Public Health; TW005572 [T.R.B., and R.R.]) and Emerging Global Leader Award; K43 TW010362 [T.R.B.]). The funders had no role in study design, data collection and analysis, decision to publish, or preparation of the manuscript. We are also grateful to the Governments of Bangladesh, Canada, Sweden and the UK for providing core/unrestricted support.

## Author contribution statements

Conceived and designed the experiments: TRB, AW, RR, FQ; Performed the experiments: RR, AW, TRB, RH, LH, AP; Analyzed the data: RR, TRB; Contributed reagents/materials/analysis tools: FQ SBC, ETR, JBH; Wrote the paper: RR, TRB, FQ, SBC, ETR, AW, JBH; Enrollment of volunteers and clinical evaluation: AIK, FC.

## Conflict of interest statement

None of the authors report any conflict of interest for the work reported here.

